# Methods for Extraction and Detection of Bacteriophage DNA from the Sputum of Patients with Cystic Fibrosis

**DOI:** 10.1101/2020.03.10.986638

**Authors:** Elizabeth B Burgener, Patrick R Secor, Michael C Tracy, Johanna M Sweere, Elisabeth M Bik, Carlos E Milla, Paul L Bollyky

**Affiliations:** Center for Excellence in Pulmonary Biology, Department of Pediatrics, Stanford University, Stanford, CA; Division of Infectious Diseases and Geographic Medicine, Department of Medicine, Stanford University, Stanford, CA; Division of Biological Sciences, University of Montana, Missoula, MT; Harbers Bik LLC, San Francisco, CA

**Author notes:** Co-senior authors (Carlos E Milla, Paul L Bollyky).

## Abstract

There is increasing interest in the pulmonary microbiome’s bacterial and viral communities, particularly in the context of chronic airway infections in cystic fibrosis (CF). However, the isolation of microbial DNA from the sputum from patients with CF is technically challenging and the optimal protocols for the analysis of viral species, including bacteriophage, from clinical samples remains challenging. Here, we evaluate a set of methods developed for processing and analyzing sputum from patients with CF with a particular emphasis on detecting bacteriophage viron-derived nucleic acid. We evaluate the impact of bead-beating, deoxyribonuclease digestion, and heating steps in these protocols focusing on the quantitative assessment of *Pseudomonas aeruginosa* and Pf bacteriophage in sputum as a proof of concept. Based on these comparative data, we describe an optimized protocol for processing sputum from patients with CF and isolating DNA for PCR or sequencing-based studies. These studies will facilitate future assessments of bacteriophage and bacteria in sputum from patients with CF.

**Author Summary:** Patients with cystic fibrosis (CF) have thickened secretions and develop chronic infections in their airways. While we culture bacterial pathogens from expectorated sputum this method favors detection of certain organisms. There is greater understanding that the microbiome with in the airway of patients with cystic fibrosis is varied and contains more than just the bacteria that we selectively culture. Processing sputum is difficult as it is very thick. There are many different ways to process sputum depending on what aspect of the sputum is to be studied. We have an interest in a specific bacteriophage, or a virus that infects bacteria. We sought to find the best method to take sputum from a CF patient and extract DNA so that we could detect both bacteria and bacteriophage DNA. We compared different methods that included different combinations of heat, mechanical homogenization, chemical lysis and DNA extraction by commercially available kits. We describe the method we found easiest to execute and produced the best yield for detection of both bacteria and bacteriophage. Our purpose of publishing this method in detail is to facilitate further study of viral and bacterial communities in the sputum of patients with CF.

## Introduction

Cystic fibrosis (CF) is an autosomal recessive, lethal disease characterized by defective anion transport across mucosal surfaces.[1] In the airways, this defect leads to thickened secretions, chronic infection, and persistent inflammation. Over time, this results in destruction of lung tissue and chronic obstructive pulmonary disease.[2] While other organ systems are affected, the aforementioned pulmonary sequelae are the major cause of morbidity and most common cause of mortality in patients with CF. While there have been substantial advancements in the treatment of airway infections and the promotion of airway clearance, microbial pathogens remain a substantial problem in CF. The study of microbial pathogens within sputum from patients with CF is therefore crucial to understanding the pathogenesis of lung infections in CF.

Recently, there is growing interest in the viruses of bacteria known as bacteriophage or simply “phage”. Due to increasing rates of antibiotic resistance, there is motivation in developing lytic bacteriophage as treatments for chronic bacterial infections to supplement or perhaps even replace conventional antibiotics. Phage therapy has a longstanding history in Eastern Europe and is making a resurgence in the West as well.[3] Following on some success in other settings, case reports of patients with CF successfully treated with phage suggest phage therapy is a viable option that deserves further investigation.[4–7] While many of these efforts have focused on *Pseudomonas aeruginosa(P. aeruginosa)* due to its high level of antibiotic resistance[4,8–10], there may be broader applications for this approach beyond treatment of *P. aeruginosa*.[5,6,11–13] As part of this development of phage for therapeutic use there is also a need to understand how phage impact mammalian immunity.[14–16]

While interest in phage is primarily related to their potential role in controlling infection, many of the bacteria that infect and colonize patients carry and express endogenous phage. Further, there is also a growing appreciation that phage profoundly shape the ecology of the lung microbiome.[17–19] Our research has focused on one such phage, Pf, a lysogenic filamentous phage that infects a portion of clinical *P. aeruginosa* isolates, is incorporated into its genome as a prophage, and is capable of replicating without killing or lysing its bacterial host. We and others have recently reported that Pf phage influence the pathogenesis of *P. aeruginosa* infections in the lung and other environments by inducing liquid crystal biofilm and promoting chronic infection.[11,12,15–17,20–22] For all of these reasons there is a pressing need for effective and reliable protocols for assessments of both phage and bacteria in CF sputum.[23,24]

Respiratory secretions are primarily composed of polymeric mucins (MUC5AC and MUC5B) that form a gel like material with complex rheological properties that form the basis for its function as an innate defense barrier, including viscoelasticity, cohesiveness, adhesiveness and wetness.[25] However, CF respiratory secretions not only have abnormal rheological properties but as a consequence of chronic infection and inflammation contain increased amounts of actin, DNA, serum proteins, lipids and bacterial-derived alginate.[26] Therefore, it is more appropriate to refer to CF expectorated secretions as sputum and not mucus.[27–29] Additionally, in CF sputum there is a greater abundance of inflammatory cells. These cells tend to necrose and provide the abundant DNA found in the CF sputum.[30] Given its multiple components and rheological properties, it is challenging to work with CF sputum. Its complex composition also renders it highly heterogenous making it quite difficult to process. Finally, CF sputum can be quite variable in its properties both between and within patients, as well as disease states, which hampers the ability to conduct comparisons between studies. This stresses the importance of establishing rigorous standardized protocols.

There are several reported methods for processing sputum from patients with CF for both detection of microbes and for measurement of proteins, cytokines and other components.[31–43] Most of these typically include a combination of steps for chemical lysis, mechanical lysis, dilution, heating, fractionization, and the addition of protease inhibitors. Most are usually tailored to the intended downstream application, including the sputum fraction that is of interest. For example, for studies of sputum inflammatory markers, such as cytokines or elastase, the sputum supernatant or matrix fraction (with the cell fraction removed) is the most relevant. In contrast, for studies of sputum bacterial components the cellular fraction is of interest. For studies of sputum mechanical or physical properties homogenization or lysis would be left out as these steps might affect those results.

In studies evaluating the presence and/or quantity of specific pathogens by PCR, similar methods are reported with chemical lysis[34,44–46], and mechanical lysis (sonication)[34,45,47], followed by DNA extraction though only involving the cellular fraction[45,46]. While some reports include detailed descriptions of methods, many unfortunately do not provide detailed protocols. For our specific area of interest, optimal protocols for isolating bacteriophage from CF sputum are unclear. We identified four studies where bacteriophage detection was reported from sputum, and they all involved dilution, chemical lysis and filtration to remove the cellular components.[48–51] Further steps to purify the phage or viral component of the sputum are taken using phenol/chloroform[50] or cesium chloride[51].

We have sought to provide a resource on the optimal methods for the collection and processing of sputum samples from patients with CF for microbial DNA studies, with a particular emphasis on phage. To this end, our protocols have been optimized for collecting DNA from Pf bacteriophage and the bacterium which produces it, *P. aeruginosa*.[20] We have focused on a DNA phage here, though it would be straightforward to modify this protocol to examine RNA viruses. These protocols reflect our recent experience in quantification and sequencing of these microbes.[15–17,20] To our knowledge this is the first systematic approach to develop a standardized method that could be used by other investigators and thus allow for meaningful comparisons. Our hope is that sharing our optimized protocols will facilitate future studies on both bacteria and phage in CF airway infections.

## Results

### DNA extraction is necessary for detection of Pf phage

We initially trialed detection of Pf phage in sputum supernatant prior to working with neat sputum. Sputum obtained from a patient with CF not infected with *P. aeruginosa* (patient CF1) was divided into two aliquots. Neat sputum was simply diluted with PBS as our group has reported previously[52] and spun to remove mucus and cells, creating a sputum supernatant. Purified Pf phage virions at a concentration of 5 × 10^10^ copies/mL were added to the sputum supernatant to assess both the detection of Pf phage in sputum and determine if DNA extraction is necessary. Both aliquots were subsequently divided into two aliquots, one of which underwent heating and DNA extraction by commercial extraction kit for genomic DNA extraction, and the other had no further processing prior to performance of qPCR (table 2) for Pf phage detection and quantification. Only the sputum supernatant which underwent DNA extraction demonstrated detectable Pf phage by qPCR (Figure 1), indicating that DNA extraction is a necessary step for releasing Pf phage DNA.

**Figure 1:**
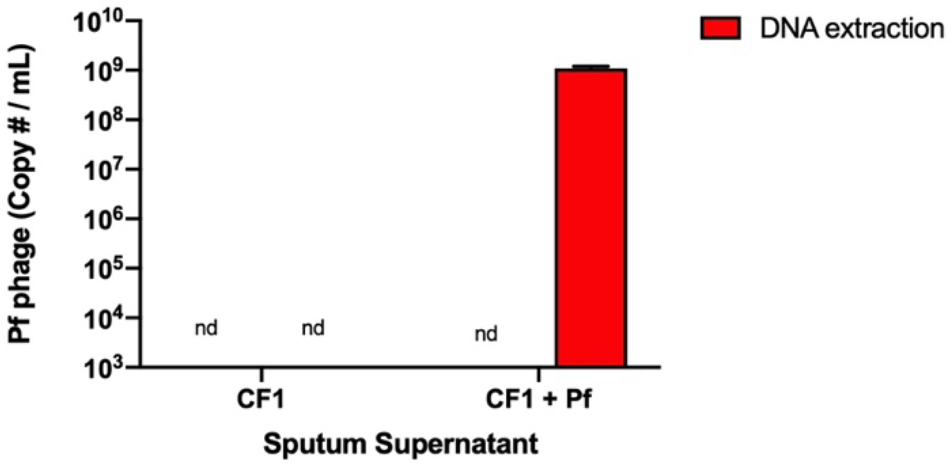
DNA extraction is necessary to detect Pf phage in sputum. Pf phage (5 × 10^10^ PFU/mL) was added to a sputum supernatant aliquot from patient CF1. Each sputum supernatant aliquot was divided into 2 further aliquots. No further processing was performed to one aliquot prior to qPCR (first bar). For the remaining aliquot, DNA extraction was performed prior to qPCR (second bar). Data are representative of three experiments with sputum from two donors. nd = not detected

### Pf phage can be detected in CF sputum

We next wanted to evaluate the detection of Pf phage in homogenized sputum (which includes cells and mucins as opposed to supernatant), to evaluate for presence of qPCR inhibitors in sputum. We added exogenous purified Pf phage at a concentration of 5 × 10^10^ copies/mL to unprocessed sputum from a patient not infected with *P. aeruginosa* (patient CF2). A second aliquot of sputum from the same patient (CF2) did not have phage added. Both were divided into two aliquots for processing.

As sputum from CF patients is thick it must be dissociated prior to DNA extraction. To optimize the DNA extraction from sputum for both the detection and quantification of Pf phage and *P. aeruginosa*, we evaluated two different methods of sputum dissociation. The first utilizes a combination of dilution, chemical lysis with DNase and dithiothreitol, heat, and followed by use of a commercially available DNA extraction kit designed for isolation of plasmid DNA[17]. We from here on refer to this method as *chemical dissociation*. The second method includes a combination of mechanical homogenization with bead beating and heat followed by use of a commercially available DNA extraction kit designed for isolation of genomic DNA.[20] We refer to this method as *mechanical dissociation.* After processing Pf phage were measured in all aliquots using qPCR (Figure 2) with significantly more Pf phage detected in the sample processed by mechanical dissociation. In concordance with patients’ respiratory culture results no *P. aeruginosa* was detected by qPCR in any of the samples.

**Figure 2:**
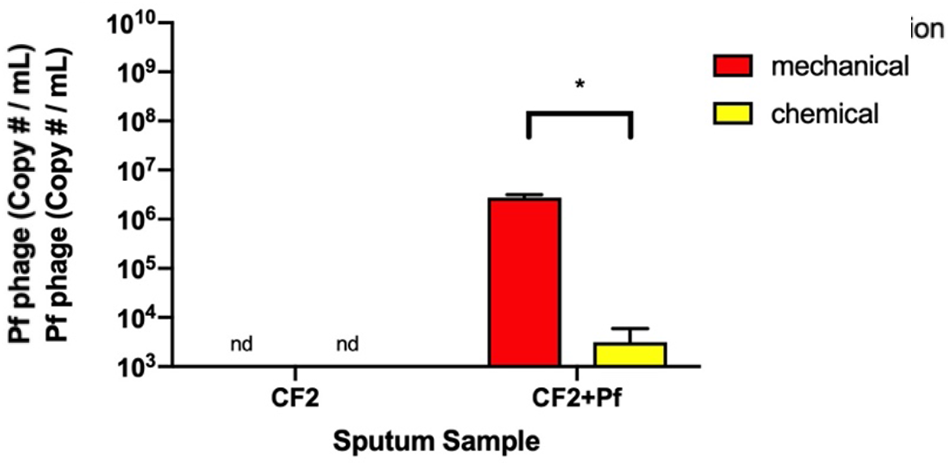
Pf phage is detected when added to CF sputum at higher levels with mechanical dissociation. Pf phage virions (5 × 10^10^ PFU/mL) were added to an aliquot of unprocessed sputum from a patient not infected by *P. aeruginosa* (patient CF2). DNA extraction with mechanical dissociation and chemical dissociation was performed prior to qPCR. Data are representative of three experiments with sputum from two donors. * =p<0.05 via a student’s T-test, nd = not detected

When Pf phage was added to both unprocessed sputum and sputum supernatant, Pf phage was detectable by qPCR at the level of 10^8^-10^9^ copies per mL sputum, demonstrating it is possible to detect Pf phage present in CF sputum with our qPCR protocol (Figures 1 & 2). While phage was detected, it was 1-2 logs lower than expected indicating that some amount of phage DNA is lost in the sputum homogenization, processing and DNA extraction.

### Mechanical dissociation improves detection of both *P. aeruginosa* and Pf phage in CF sputum

We next then evaluated the presence of Pf phage and *P. aeruginosa* in the sputum of patients with CF with known *P. aeruginosa* infection. Sputum was aliquoted and processed by both mechanical and chemical dissociation procedures followed by quantification of *P. aeruginosa* and Pf phage by qPCR(Figure 3A) and quantification of total nucleic acid content(Figure 3B). Both chemical and mechanical dissociation procedures resulted in detectable *P. aeruginosa* and Pf phage in the patient sputum with history of *P. aeruginosa* infection, however the mechanical dissociation appears to preserve the yield of detection as well as increase nucleic acid yield.

**Figure 3:**
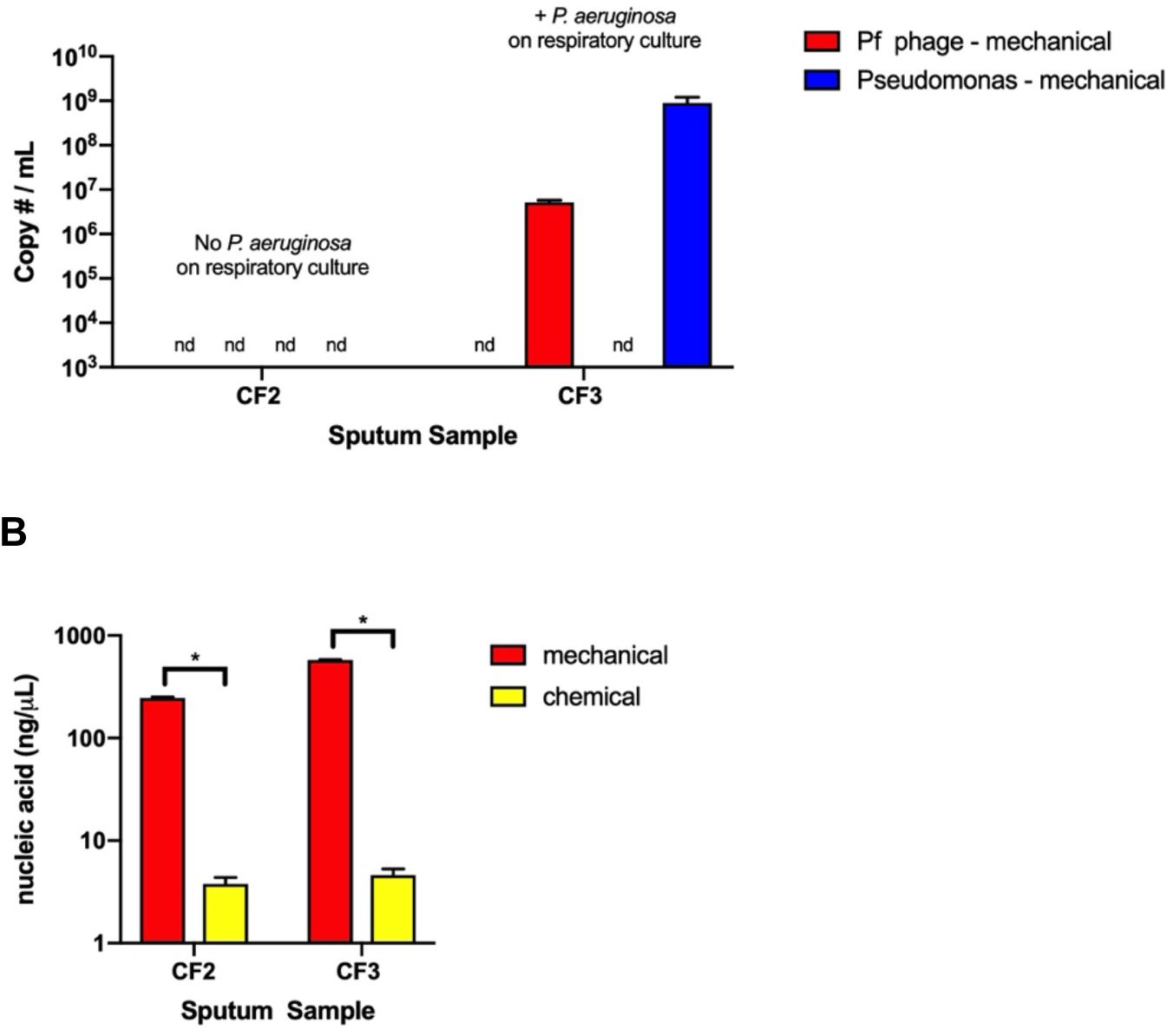
Comparison of dissociation methods for DNA extraction. We directly compared DNA extraction using the chemical dissociation method [17] and mechanical dissociation method[20] in a sputum sample from a patient that was negative for *P. aeruginosa* by respiratory culture (patient CF2) and in a patient who does grow *P. aeruginosa* on respiratory culture (patient CF3). The sample from each patient was divided into 2 aliquots and one dissociated by each method. Results are show in the following order: chemical-Pf phage, mechanical-Pf phage, chemical - *P. aeruginosa*, mechanical – *P. aeruginosa* **A)** *P. aeruginosa* and Pf phage copies detected by PCR. **B)** Nucleic acid content after DNA extraction. Data is representative results of two replicate experiments with sputum from two donors. * =p<0.05 via a student’s T-test, nd = not detected

### Pf phage can be detected over a range of concentrations

To ensure appropriate detection of Pf phage by qPCR, we examined the performance of our protocol over a range of Pf concentrations. We observed in a sample with Pf phage added to the sputum that dilution of the sample leads to quantification that corresponded to the dilution factor (Figure 4). These data indicate that by using this protocol Pf phage can be detected over a range of concentrations.

**Figure 4:**
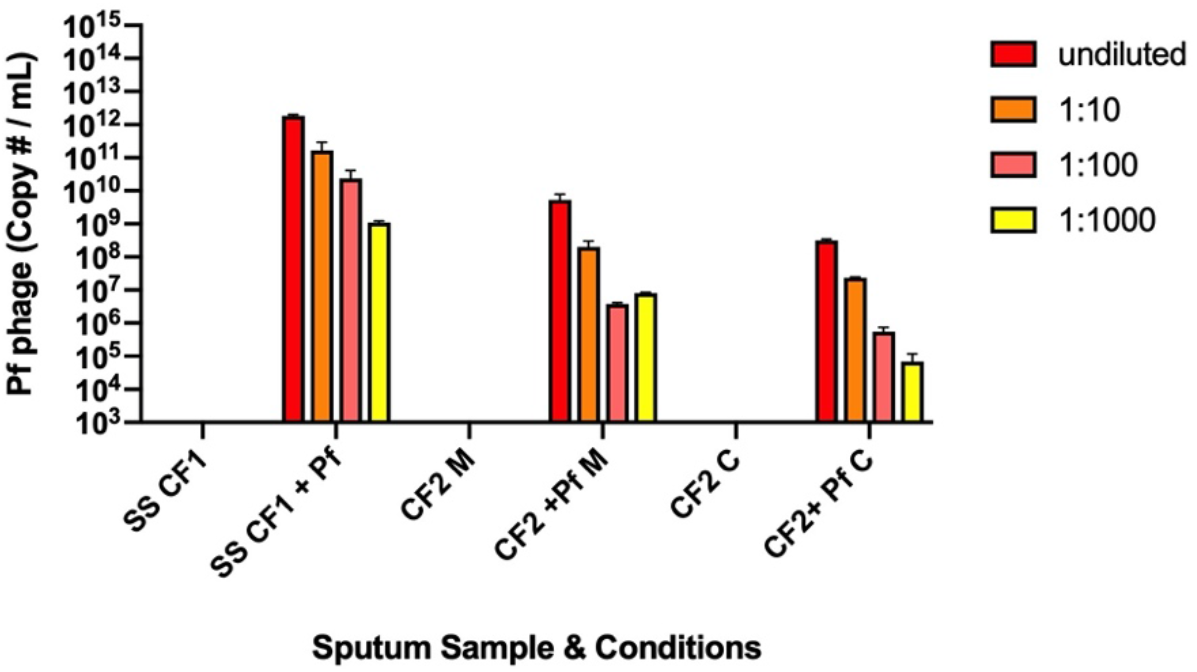
Pf phage are detectable over a range of concentrations. Pf phage quantified by PCR in dilutions of the DNA extracts from the samples from Fig. 1, 2 and 3. M= mechanical dissociation, C = chemical dissociation, SS = sputum supernatant

## Discussion

We set out to establish a protocol that would allow for optimal detection and quantification of both phage and bacteria by PCR in the sputum of patients with CF. Different processing conditions were investigated by combining homogenization, lysis, heat, fractionation, and DNA extraction. We identified a protocol which utilizes mechanical dissociation, heat and DNA extraction by commercially available kit as the method that gave the best performance in its quantification of both Pf phage and *P. aeruginosa* in CF sputum. We also found that sputum supernatant by itself did not interfere with the detection of Pf phage. This experiment also demonstrated that heating in addition to a DNA extraction method is necessary, which is not surprising given the Pf phage coat protein is very stable up to a temperature of 70°C[53] and heating likely facilitates the release of Pf phage DNA. Given that Pf genes are integrated into the *P. aeruginosa* chromosome, this is of importance as it allows for the detection of not only Pf prophage but also the free or extracellular phage present in the sample.

While we trialed 2 DNA extraction methods both from commercially available kits, we found the kit designed for genomic DNA isolation yielded more consistent results than the kit designed for isolation of plasmids. The kit designed for plasmid detection/recovery will capture small DNA segments of <10 kilo base pairs (kbp) in length.[54] The genomic DNA kit which yielded better DNA recovery and more consistent results for both Pf phage and *P. aeruginosa* is designed for the isolation of genomic, bacterial, mitochondrial and viral DNA and reported to recover DNA segments of approximately 20-30 kilo base pairs in length.[55] We suspected we would have better Pf phage recovery with the Miniprep kit because the Pf phage genome is approximately 5-12 kbp in length, depending on strain[56–58], however this was not what we observed. This is likely a result of the centrifugation step prior to DNA extraction, as Pf phage may be pulled down with the cells and bacteria in the sputum, rather than a difference in the DNA extraction kits.

A limitation of this study is the lack of a second confirmatory testing for quantification of Pf phage in the sputum samples assayed. As Pf phage is lysogenic (not a lytic phage) it is difficult to use plaque assays to quantify Pf phage. Lysis can be seen when Pf phage production is high or from a purified phage preparation and we have previously quantified phage and validated our Pf phage PCR.[17] However, a plaque assay performed on CF sputum would not confirm presence of Pf phage as there are too many other substances, microbes and possibly other bacteriophage present in CF sputum that could cause plaque formation. Given plaque assays were unsuccessful and would be non-specific for Pf phage, we circumvented this limitation by spiking known concentrations of exogenous Pf phage into the sputum samples and demonstrated their detection with qPCR.

In conclusion, we describe our experiments that led to our protocols (Table 1, 2 and Figure 5) for extraction and detection of phage and bacterial DNA by mechanical dissociation followed by qPCR from the sputum of patients with CF. We demonstrate the performance characteristics of our method by testing both Pf phage and *P. aeruginosa* presence and quantity by qPCR. While there are multiple reported methods for the processing and analysis of CF sputum, we found the use of whole sputum processed with mechanical dissociation followed by heating and DNA extraction proved to have the highest and most consistent nucleic acid yield for detection of both viral and bacterial components (Figure 5).

**Table 1:** Protocol for Mechanical Dissociation and DNA Extraction as used in Burgener et al.[20].

|  |
| --- |
| Reagents:<br>• Ethanol<br>• Proteinase K and buffers from QIAmp DNA Mini Kit |
| Equipment<br>• 18 gauge needles<br>• 3mL syringes<br>• pipettes<br>• Ceramic beads (1 mm) & 2mL impact resistant sterile tubes with screw cap<br>• Mechanical homogenizer such as MagNA Lyser Instrument, Roche<br>• Dry block incubator<br>• 2mL spin columns from QIAmp DNA Mini Kit<br>• 1.5mL Eppendorf tubes<br>• Vortex mixer<br>• Centrifuge |
| Protocol:<br>1. Thaw sputum at room air.<br>2. If unable to pipette sputum, then use 18g needle to homogenize, 5-10 slow pumps up and down. Then pipette 200µL of sputum sample in bead beating tube with approximately 1/5 filled with ceramic beads.<br>3. Add 20µL of proteinase K and 200µL of buffer AL to the sample.<br>4. Vortex for 15 seconds<br>5. Bead beat for 30 seconds at 6500 RPM. Place in cooling rack for ~1-2 minute. Beat again for 30 seconds at 6500 RPM.<br>6. Incubate at 56°C for 10 minutes.<br>7. Incubate at 95°C for 5 minutes for lysis and disinfection.<br>8. Add 200µL ethanol to the sample in the bead beating tube.<br>9. Vortex for 15 seconds, followed by brief centrifuging of the tube.<br>10. Carefully apply mixture to QIAmp spin column in a 2mL collection tube. Centrifuge at 6000xg (8000 rpm) for 1 minute.<br>11. Place the column in a clean 2mL collection tube. Pour the flow through into a toxic waste bottle (buffer AL contains guanidine hydrochloride), and discard the empty collection tube.<br>12. Add 500 µL wash buffer AW1 to spin column. Centrifuge at 6000xg for 1 min.<br>13. Place column in a clean 2mL collection tube (kit). Pour the flow through into a toxic waste bottle (buffer AW1 contains guanidine hydrochloride), and discard the empty collection tube.<br>14. Add 500 µL wash buffer AW2. Centrifuge at 20,000xg (or 14,000rpm) for 3 minutes.<br>15. Place the column in a clean 1.5mL Eppendorf tube (not provided by kit) and discard the tube containing the flow through (buffer AW2 does not contain guanidine hydrochloride).<br>16. Add 200µL elution buffer AE. Incubate at room temperature for 5 minutes.<br>17. Centrifuge at 6000g (8000 rpm) for 1 minute.<br>18. Store DNA at -20C or -80°C. |

**Table 2:** Protocol of qPCR for Pf phage and *Pseudomonas aeruginosa* used in Burgener et al and Sweere et al.*[15,20]*.

|  |
| --- |
| Materials:<br>• Taqman SensiFAST™ Probe Hi-ROX (Bioline, Cat. No. BIO-82020)<br>• Forward primer, specific for *rpIU* or *Pfcon*<br>• Reverse primer, specific for *rpIU* or *Pfcon*<br>• Probe, specific for *rpIU* or *Pfcon*<br>• RNase free water |
| Equipment:<br>• Cooling block or ice and ice bucket<br>• MicroAmp Fast Optical 96 well plates<br>• StepOne Plus Real Time PCR System |
| **Protocol**<br>Prepare “master mix”<br>1. Prepare a total of 9µL for each well + 1µL sample (samples run in triplicate, water as negative control run in triplicate and 21 wells for standard)<br>a. Taqman 5µL (SemiFAST Probe HI-Rox Mix)<br>b. Forward primer 1µL, specific for *rpIU* or *Pfcon*<br>c. Reverse primer 1µL, specific for *rpIU* or *Pfcon*<br>d. Probe 1µL, specific for *rpIU* or *Pfcon*<br>e. RNase free water 1µL<br>2. Keep master mix on ice.<br>Fill plate with master mix followed by samples:<br>1. Use 96 well plate (MicroAmp Fast Optical) placed in cooling block or ice<br>2. Fill each well with 9µL with corresponding master mix, *rpIU* or *Pfcon*.<br>3. Add 1 µL of samples and water/negatives to appropriate wells.<br>4. To minimize contamination of samples cut plate cover to cover samples and water wells. Add plate cover, pressing down with Kimwipe (not hands).<br>5.<br>Prepare standard dilution ladder and add to plate:<br>1. Prepare plasmid standard (50ng/µL) of both *Pfcon* and *rpIU* in 1:10, 1:100, 1:1,000, 1:10,000, 1:100,000, 1:1,000,000 and 1:10,000,000<br>2. Plate standard dilution in triplicate with 1µL of standard (in place of sample).<br>3. Use remainder of plate cover to cover remaining wells.<br>4. Centrifuge plate – pulse for ~30 sec, >12G. This helps remove bubbles and seal cover.<br>Run PCR on StepOne Plus Real Time PCR System:<br>1. Settings: 96 wells, quantitation with standard curve.<br>2. Select Taqman reagents and “Fast” protocol<br>3. Input location of samples and standard dilution in plate plan.<br>4. In set up screen:<br>a. Set well volume to 10µL<br>b. Holding Stage. Set Step 1 to 95°C and 2:00<br>c. Cycling Stage - Step 1 to 95°C and 00:15. Step 2 to 60°C and 00:20<br>d. Set for 40 cycles |

**Figure 5:**
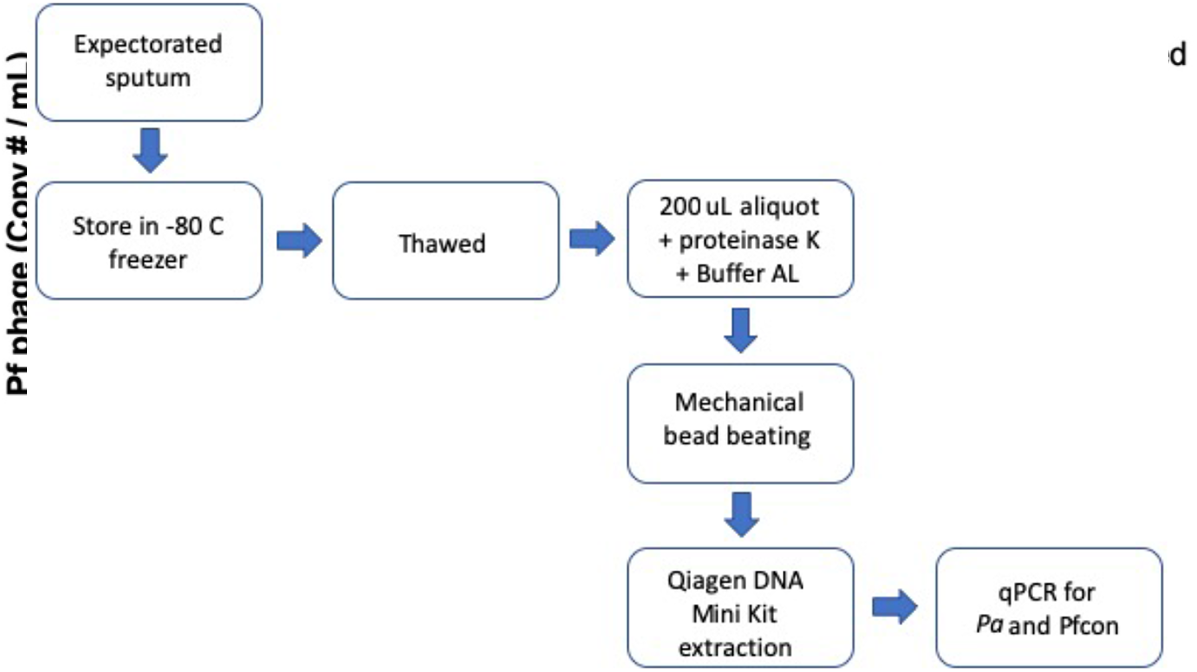
Flow chart for sputum processing used in Burgener et al. Starting with collection of expectorated sputum collected in clinic followed by freezer storage, DNA extraction and performance of qPCR for both Pf phage and *Pseudomonas aeruginosa (Pa)*.

## Methods

### Ethics Statement

Approval for sputum collection was obtained via the Stanford University Institutional Review Board (IRB #11197). All patients signed written informed consent documentation.

### Patient Selection

Patients were recruited from the Stanford University Cystic Fibrosis Center. For optimization of sputum processing inclusion criteria was a confirmed diagnosis of CF and exclusion criteria was lung transplantation. Three patients were identified in clinic visits or during hospitalization and had conventional testing, respiratory culture, as per standard CF clinical care to test for the presence of *P. aeruginosa*.

### Collection and Storage of Sputum

Expectorated sputum was collected from subjects in sterile specimen cups and frozen at −80°C for later batch processing. We chose to use expectorated sputum only as we felt throat swabs, induced sputum and bronchoalveolar lavage specimen would not yield comparable concentrations of bacteria and bacteriophage if we included all specimen types. Expectorated sputum was the most available and easily collectable specimen type.

### Processing of Neat Sputum to Sputum Supernatant

Sputum was thawed at room temperature prior to processing. The sample was weighed and then diluted 1:9 by weight with PBS. Agitation was performed with 18 gauge needle and 3mL syringe by pumping up and down for 20 strokes as our group has previously reported.[52] The homogenized sample was passed through a 70um cell filter and the filtered homogenate was centrifuged at 400g for 10 minutes at 4°C to separate the cell fraction. The supernatant was carefully decanted to a new tube and centrifuged at 3000g for 10 minutes at 4°C. The resulting supernatant was stored at −80°C in 200uL aliquots for later DNA extraction. The sample was then used to perform qPCR as described below.

### Addition of Exogenous Pf Phage

1uL of purified Pf phage preparation (1 × 10^13^ PFU/mL) was added to 200μL of sputum supernatant or sputum sample yielding a concentration of 5 × 10^10^ copies/mL in the sample. Pf phage purification and preparation is described in our previous manuscript Sweere et al.[15] Briefly, Pf phage virions were precipitated with NaCl and polyethylene glycol (MW 8000) from the filtered supernatants of infected *P. aeruginosa* cultures. Phage pellets were then dialyzed to remove residual salts and polyethylene glycol and then titered.

### Chemical Dissociation and DNA Extraction of Neat Sputum

This protocol was used by Secor et al. for detection of Pf phage and *P. aeruginosa*.[17] Neat sputum was diluted 1:1 with PBS and DNase (100ug/mL) and DTT (1mM). The homogenate was then incubated at 37°C for 5 hours with occasional vortexing. The sample was then centrifuged at 6,000G for 10 minutes and the supernatant collected. A 100uL aliquot was incubated at 95°C for 15 minutes to inactivate DNAse and release bacteriophage DNA from its protein coat. The sample was then added to 250μL of buffer P1 from QIAGEN MiniPrep Kit. The DNA extraction was then performed as per manufacturers protocol.

### Mechanical Dissociation and DNA Extraction of Neat Sputum

This protocol was designed for microbiome analysis in sputum and was adapted and used in our recent manuscript Burgener et al.[20] Briefly, if necessary for aliquotting, neat sputum was crudely homogenization with 5-10 pumps of 18g needle and syringe. Approximately 200uL of neat sputum was added to a 2.0mL tube filled approximately 1/5 by volume with 1 mm ceramic beads. Proteinase K (20 uL) and buffer AL (200 uL) from the QIAamp DNA Mini Kit by QIAGEN was added prior to the samples undergoing mechanical bead beating (MagNA Lyser Instrument, Roche) for 60 seconds at 6500 rpm followed by 2-3 minutes of cooling and a second round of 60 seconds at 6500 rpm. The homogenate was then incubated for lysis and disinfection. DNA extraction was then performed as per the manufacturer’s protocol for tissue starting with the steps after buffer AL addition. DNA was eluted by adding 200 uL of elution buffer AE. This is described in detail in Table 1.

### Nucleic Acid Measurement

Yield of nucleic acid after DNA extraction was measured using a NanoDrop 2000 Spectrophotometer (Thermo Fisher Scientific) in 2uL of sample.

### Dilution of DNA Extracted from Sputum Samples

DNA extracts from sputum were diluted with RNAse free water 1:10, 1:100 and 1:1000.

### Pf Phage and *P. aeruginosa* PCR

Pf phage and *P. aeruginosa* in sputum was quantified by quantitative PCR as we have previously described.[15,20] Briefly, 2uL of the DNA extracted from sputum samples was used as a template in 20-mL qPCR reactions containing 1× SensiFAST™ Probe Hi-ROX (Bioline, Cat. No. BIO-82020), 200 nM probe, and 2 nM forward and reverse primers. Cycling conditions were as follows: 95°C for 2 min, (95°C for 15 sec., 60°C for 20 sec.) × 40 cycles on a StepOnePlus Real-Time PCR system (Applied Biosystems). To quantify Pf, the primers (Pf-Conserve-F (TTCCCGCGTGGAATGC) and Pf-Conserve-R (CGGGAAGACAGCCACCAA)) targeting Pfcon (*PA0717*), a gene that is conserved across diverse strains of Pf phage (Pf1, Pf4, Pf5), were used together with a PA0717 probe (AACGCTGGGTCGAAG). For *P. aeruginosa* quantification, primers targeting the 50S ribosomal protein gene *rpIU* (rpIU-F (CAAGGTCCGCATCATCAAGTT) and rpIU-R (GGCCCTGACGCTTCATGT)) were used together with a *rpIU* probe (CGCCGTCGTAAGC). For the standard curves, the sequence targeted by the primers and probe were inserted into a pUC57 plasmid (Genewiz) and tenfold serial dilutions of the plasmid were used in the qPCR reactions. Using these methods, we determined that the lower limit of detection in our qPCR assay for the *rpIU* gene of *P. aeruginosa* is 10^2^ copies per mL and for the Pf phage conserved gene is 10^3^ copies per mL. All samples were run in duplicate or triplicate. The reported Pf phage concentrations represent the measured Pf phage with the *P. aeruginosa* concentration subtracted to account for detection of prophage DNA contained in the *P. aeruginosa* genome. Protocol steps are described in Table 2. This protocol was used in both of our manuscripts Sweere et al. and Burgener et al.[15,20]

## Acknowledgements

We thank Alexander Yacob and Daniel Bassily for assistance in processing sputum samples and experimental execution.

